# A Chromosome-length Assembly of the Black Petaltail (*Tanypteryx hageni*) Dragonfly

**DOI:** 10.1101/2022.10.18.512723

**Authors:** Ethan R. Tolman, Christopher D. Beatty, Jonas Bush, Manpreet Kohli, Carlos M. Moreno, Jessica L. Ware, K. Scott Weber, Ruqayya Khan, Chirag Maheshwari, David Weisz, Olga Dudchenko, Erez Lieberman Aiden, Paul B. Frandsen

## Abstract

We present a chromosome-length genome assembly and annotation of the Black Petaltail dragonfly (*Tanypteryx hageni*). This habitat specialist diverged from its sister species over 70 million years ago, and separated from the most closely related Odonata with a reference genome 150 million years ago. Using PacBio HiFi reads and Hi-C data for scaffolding we produce one of the most high quality Odonata genomes to date. A scaffold N50 of 206.6 Mb and a BUSCO score of 96.8% indicate high contiguity and completeness.

**Significance:** We provide a chromosome-length assembly of the Black Petaltail dragonfly (*Tanypteryx hageni)*, the first genome assembly for any non-libelluloid dragonfly. The Black Petaltail diverged from its sister species over 70 million years ago. *T. hageni*, like its confamilials, occupies fen habitats in its nymphal stage, a life history uncommon in the vast majority of dragonflies. We hope that the availability of this assembly will facilitate research on *T. hageni* and other petaltail species, to better understand their ecology and support conservation efforts.

## Introduction

The Black Petaltail dragonfly (*Tanypteryx hageni*), found in montane habitats from California to British Columbia, is something of an evolutionary enigma. It is a member of the odonate family Petaluridae (known as ‘petaltails’ due to the broad, petal-like claspers at the end of the male abdomen), which is estimated to have originated approximately 150 million years ago (Ware et al. 2014)(fig. 1). The relative position of Petaluridae with respect to other dragonfly (suborder Anisoptera) families has varied with taxon sampling, data source, and phylogenetic reconstruction method (e.g., (Suvorov et al. 2021; Bybee et al. 2008; Letsch 2007; Kohli et al. 2021; Blanke et al. 2013)).

**Figure 1:**
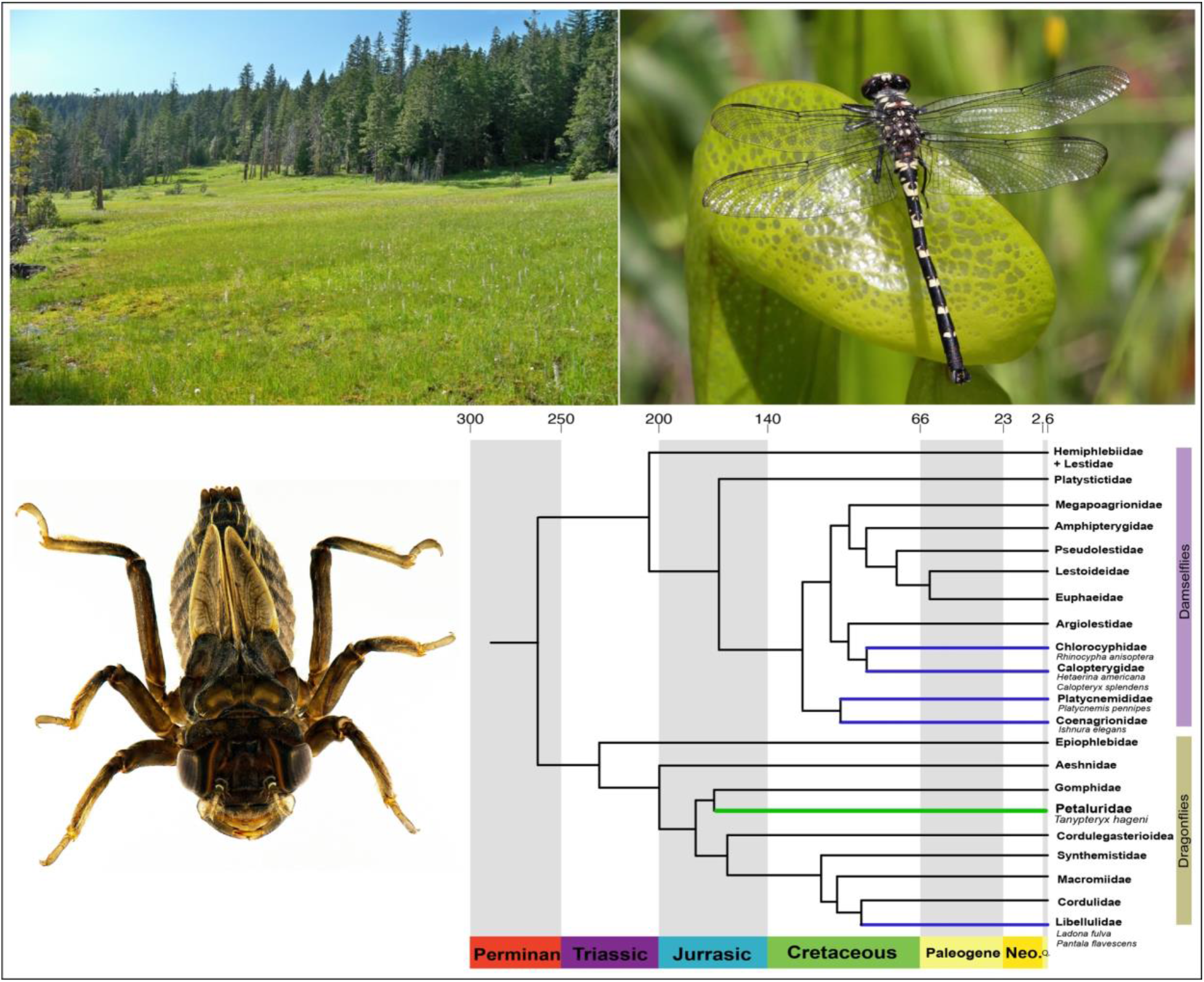
(Upper left) A fen in Lassen National Forest (California, USA) where T. hageni was collected. (Upper right) T. hageni, adult. (Bottom Left) T. hageni, larvae, credit Marla Garrison. (Bottom right) Phylogeny of Odonata (modified from Kohli et al 202).1 Families with a reference genome highlighted in purple, Petaluridae highlighted in green.

While the family originated long before most recognized insect families, the geologic ages of its member species are even more extreme in relation with other animal species, leading members of Petaluridae to be considered as “living fossils.” Most extant petaltails are estimated to have appeared in the mid- to late-Cretaceous, with speciation being driven by major events like continental drift; *T. hageni* is estimated to have diverged from the sister species *Tanypteryx pryeri* (found in Japan) ∼73 million years ago, potentially diverging when the Beringian land bridge disappeared in the late Cretaceous (Ware et al. 2014; Fiorillo 2008). Often, species with long geological persistence have wide geographic ranges (Hopkins 2011; Powell 2007), and tend to be habitat generalists, or give rise to habitat generalists (Colles et al. 2009), but for *T. hageni* this is not the case—the persistence of this species (as well as other petaltails) is puzzling as the nymphs of *T. hageni* exclusively inhabit fens (Baird 2012), groundwater-driven habitats which host a number of specialist animals and plants. These habitats are characterized by soils saturated by groundwater, commonly found around springs and in riparian areas of headwater streams. Nymphs (fig. 1) dig and maintain a burrow (a behavior displayed by a number of other petaltail species) that fills with water.

The research on fens in North America is sparse, but it is known that while fens make up a tiny fraction of the North American landscape, they contain a surprising proportion of the continent’s biodiversity. The US Department of Agriculture observes that between 15 and 20% of the rare and uncommon plant species found in Deschutes National Forest (Oregon, USA) are found in fen ecosystems (US Department of Agriculture). It is estimated that fens are the most floristically diverse wetlands in the United States, and contain a high number of rare and endangered species (Bedford & Godwin 2003). Research on fens across the range of *T. hageni* is minimal; it is known that montane fens in Oregon (a portion of *T. hageni*’s range) only occur in “low-permeability glacial-till…around 1400–1800 m in elevation, and are concentrated in areas mantled by pumice deposits that originated primarily from the eruption of Mt. Mazama approximately 7700 years BP” (Aldous et al. 2015). These Oregon fens are supplied by perched aquifers in glacial till, and are therefore unaffected by the draining of deeper regional aquifers, but they are especially susceptible to changes in recharge due to climate change (Aldous et al. 2015). It has been hypothesized that Oregon fens could be negatively affected by fire suppression (Tolman 2007), but there is scant research evaluating how recent megafires throughout the range of *T. hageni* may be influencing fens in this range. However, it is known that fens are degrading across the continental United States (Bedford & Godwin 2003). Thus, we have concerns not only for the survival of the Black Petaltail, but for the specialized habitats in which they live. There is little research regarding genomic adaptations to life in fens, so it is paramount to establish a baseline of understanding for this declining habitat.

Here, we present a chromosome-length assembly of the Black Petaltail. This genome will be a valuable tool for studying an organism that may be especially hard hit by climate change and habitat destruction. Additionally, this genome will shed light on an evolutionary enigma: the petaltail dragonflies have persisted for tens of millions of years, despite exclusively occupying fragile fen habitat as nymphs (Ware et al. 2014). Lastly, this genome will be an important resource in resolving the phylogeny of early divergences within Odonata, as no genome of any basal anisopteran is currently available.

## Results and Discussion

### Sequencing and Genome Size Estimation

We recovered >44.6 Gb of sequence contained in HiFi reads, generated from 730 Gbp of raw sequence in subreads from two PacBio SMRT cells. The estimated genome size using kmers from HiFi reads with GenomeScope 2.0 was 1.47 Gb with an estimated 59.9% of unique sequence (Ranallo-Benavidez et al. 2020), resulting in approximately 25x coverage (supplementary figure 1).

### Genome Assembly and QC

Our contig assembly was generated with hifiasm v.0.16.1 (Cheng et al. 2021) and submitted to NCBI to identify possible contaminants. Following the removal of two possible contaminants, the assembly was 1.69 Gb in length, contained 2,133 contigs, and had a contig N50 > 4 Mbp (supplementary table 1). After scaffolding with Hi-C data, we generated a highly contiguous assembly that was 1.70 gb in length with a scaffold n50 > 206.6 Mbp, with 90.465% of base pairs assigned to nine chromosomes (supplementary table 1, see supplementary fig. 5 for contact map). We filtered out contigs that were assigned to proteobacteria, mollusca, cnidaria and bacteroidetes by BLAST v.2.9.0 (Camacho et al. 2009) and replaced mitochondrial contigs with the mitochondrial genome assembly, resulting in a final assembly length of 1.68 Gbp with a scaffold N50 > 206.6 Mbp (supplementary table 1) and an overall GC content of 37.98% (supplementary table 2). We recovered 96.8% of universal single copy orthologs (including 96.2% single and complete and an additional .6% fragmented) from the BUSCO (Manni et al. 2021) Insecta database indicating a high level of completeness, especially when compared to most publicly available Odonata genomes (table 1).

**Table 1:** Compares the contiguity and completeness of available Odonata genome assemblies.

| **Table I: Comparison of publicly available Odonata genomes** |  |  |  |  |  |  |
| --- | --- | --- | --- | --- | --- | --- |
|  | Suborder | Assembly level | N50 (mb) | Scaffolds | BUSCO Score of Assembly | Source |
| *Tanypteryx hageni* | Anisoptera | Chromosome | 206.6 | 1,033 | 96.8%** | Authors |
| *Ladona fulva* | Anisoptera | Contig | .06 | 9,411 | 95.7% | NCBI |
| *Pantala flavescens* | Anisoptera | Chromosome | 553 | 43 | 96.9% | (Liu et al. 2022) |
| *Ischnura elegans* | Zygoptera | Chromosome | 123.6 | 110 | 97.2% | (Price et al. 2022) |
| *Hetaerina americana* | Zygoptera | Scaffold | 86.1* | 1583* | 97.7%** | NCBI |
| *Platycnemis pennipes* | Zygoptera | Chromosome | 144.8 | 88 | 96.9%** | NCBI |
| *Rhinocypha anisoptera* | Zygoptera | Contig | .41* | 754,445* | 73.5%** | NCBI |
| *Calopteryx splendens* | Zygoptera | Contig | .42* | 8,896* | 94.7%** | NCBI |
| *Table 1: Compares the contiguity and completeness of available Odonata genome assemblies.*<br>*\*Calculated by the authors using assembly-stats (Trizna 2020)*<br>*\*\*Calculated by the authors using BUSCO (Manni et al. 2021)*<br>*All other statistics are taken from the source.* |  |  |  |  |  |  |

After blobtools analysis, 19.22% of our genome was assigned to chordata. (supplementary fig. 2). This included two of the chromosome-length scaffolds, which had nearly identical GC proportions and coverage to the other chromosome-length scaffolds (supplementary fig. 2). It is unlikely that this was due to contamination, as our Hi-C experiment assigned these to chromosomes.We hypothesized that this could be due to a lack of coverage of the lineage Petaluridae in the BLAST database. To test this, we characterized the top BLAST hits of publicly available Petaluridae transcriptomes. In the transcriptomes of *Phenes raptor* (Suvorov et al. 2021), *Tanypteryx pryeri* (Suvorov et al. 2021), and *Tanypteryx hageni* (Misof et al. 2014) 23%, 10%, and 10% of contigs had a top blast hit of chordata (supplementary fig 3). This suggests that a lack of database coverage for *Petaluridae* could be a likely explanation for this phenomenon. It appears that the genomes of *Petaluridae* have greatly diverged from what is covered in databases, leading to erroneous BLAST characterizations. We also observed this phenomenon when blasting the genome of the Atlantic Horseshoe Crab (*Limulus polyphemus*), where 9.7% of BLAST hits mapped to Chordata (supplementary fig. 4). The Atlantic Horseshoe Crab also has a highly repetitive genome, and lack of database coverage may be resulting in BLAST results outside of the phylum.

### Annotation

55.12 % of the genome was classified as repetitive using RepeatModeler2v2.0.1 (Flynn et al. 2020) and RepeatMasker v4.1.2 (Smit et al. 2013). 26.14% of the genome was classified as “unclassified repetitive elements’’, 15.73% as DNA transposons, 11.38 % as retroelements and 1.53% as rolling circles. We identified 22,261 protein coding genes. The annotated protein set contained 89.4% of BUSCO insecta genes, with 6.7% of the BUSCO genes duplicates, and another 4.7% fragmented.

#### Mitochondrial Genome

We assembled the mitochondrial genome resulting in a circular contig with a length of 16,053 bp, and a GC content of 24.62% (supplementary table 2).

## Conclusion

Here we present one of the most complete Odonata genomes to date. As the first non-libelluloid Anisopteran genome, and the first genome of an odonate habitat specialist, this assembly will be a valuable tool for understanding the biology of the Black Petaltail, resolving the phylogeny of Anisoptera, and will provide general insights into long-persisting species.

## Materials and Methods

### Specimen collection, DNA extraction, and sequencing

We collected an immature nymph live from a burrow near Cherry Hill Campground (Lassen National Forest, California, USA) in fall of 2020. The specimen was flash frozen in liquid nitrogen and stored in a −80 °C freezer prior to extraction. High molecular weight DNA was extracted from a single individual using the Qiagen Genomic-tip kit.

Another specimen was collected from the same location in spring 2022 for Hi-C library generation. It was also flash frozen in liquid nitrogen, and sent for Hi-C analysis by DNA Zoo, who used the hemolymph for Hi-C library preparation.

High molecular weight DNA was sheared to 18 kbp using a Diagenode Megaruptor and prepared into a sequencing library using the PacBio HiFi SMRTbell^®^ Express Template Kit 2.0. The library was sequenced on two PacBio Sequel II 30 hour SMRT cells in CCS mode at the BYU sequencing center.

### Sequencing QC and genome assembly

We generated HiFi reads from raw subreads with PacBio SMRTlink. We then used the HiFi reads to estimate genome size with Genomescope 2.0 and smudgeplot (Ranallo-Benavidez et al. 2020). We generated an initial contig assembly with hifiasm v.0.16.0(Cheng et al. 2021), and submitted the assembly to NCBI, through the genome submission tool, to check for contamination.

To generate chromosome length scaffolds, we used High-throughput chromosome conformation capture (Hi-C). The DNA Zoo consortium (dnazoo.org) generated an in situ Hi-C library using the protocol described in Rao et al. (Rao et al. 2014). Hi-C data was then aligned to the draft assembly using Juicer (Durand et al. 2016), and the candidate chromosome length genome assembly was built using 3D-DNA (Dudchenko et al. 2017). The resulting contact maps (supplementary fig. 5) were manually reviewed using Juicebox Assembly Tools (Durand et al. 2016; Dudchenko et al. 2018) Interactive contact maps were generated using juicebox.js (Robinson et al. 2018), for both the draft and reference assembly and are publicly available at https://www.dnazoo.org/assemblies/Tanypteryx_hageni.

We screened for contamination with taxon-annotated GC-coverage plots using BlobTools v1.1.1 (Laetsch & Blaxter 2017). We mapped all Hi-Fi reads against the final assembly using minimap2 v2.1 (Li 2018), sorted the bam file with samtools v1.13 (Danecek et al. 2021) using the command *samtools sort*, and assigned taxonomy with megablast (Shiryev et al. 2007) using the parameters: *task megablast and -e-value 1e-25*. We calculated coverage using the blobtools function *map2cov*, created the blobtools database using the command *blobdb*, and generated the blobplot with the command *blobtools plot*. After examining the blobplot we removed contigs blasting to proteobacteria, bacteroides, cnidaria and mollusca.

To investigate whether excessive megablast assignments to chordata were due to a lack of database coverage for *Petaluridae*, we used BLAST to classify the transcriptomes of *Tanypteryx hageni* (Misof et al. 2014), and other members of Petaluridae, *Tanypteryx pryeri* and *Phenes raptor* (Suvorov et al. 2021), against the Genbank nucleotide database, using the same parameters as above: *task megablast* and *-e-value 1e-25*.

### Quality Control

We generated all contiguity stats with assembly-stats (Trizna 2020). We also ran BUSCO (Manni et al. 2021) on the other publicly available Odonata genomes for comparison (Table 1) using the *Insecta* database, in genome mode with the flag *--long* to retrain BUSCO for more accurate identification of genes.

### Annotation

We first modeled and masked the repetitive elements of the scaffold and chromosome-level assemblies using RepeatModeler2 (Flynn et al. 2020). We then annotated the masked, scaffold-level assembly using MAKER v3.01.03 (Campbell et al. 2014). We ran a homology-only MAKER run using the 1kite *Tanpyeryx hageni* transcriptome (Misof et al. 2014), the transcriptomes of the Petaluridae *Tanypteryx pryeryi* and *Phenes raptor* (Suvorov et al. 2021), and the complete annotated protein sets of *Ladona fulva, Pantala flavescens*, and *Ischnura elegans*. We trained Augustus (Stanke et al. 2008, 2006) on the identified protein sets using BUSCO and the insecta dataset (Manni et al. 2021), and ran MAKER a second time to generate *ab-initio* gene predictions. We then mapped these proteins to the chromosome level assembly using miniprot v0.4 (Li 2022) and re-trained Augustus (Stanke et al. 2008, 2006) using the scaffold-level coding sequences, with 1000 base pairs surrounding each sequence as the training set. As this annotation resulted in a high number of genes, and a less-than ideal BUSCO score we also mapped the protein set of *Pantala flavescens* (Liu et al. 2022) to the masked chromosome level assembly using miniprot v0.4(Li 2022), and extracted the mapped *Pantala* proteins, the protein set from the augustus annotation, and the protein set of the mapped proteins from their respective gff files with *gffread* (Pertea 2022). We combined all three protein sets and clustered the proteins at 80* similarity with CD-HIT v4.8.1 (Fu et al. 2012). We then mapped this clustered protein set back to the chromosome-level assembly with miniprot (Li 2022), and used BLAST (Shiryev et al. 2007) to align the candidate proteins to all Arthropoda proteins available on NCBI using the parameters *-outfmt “6 sseqid pident evalue qseqid -max_target_seqs 1 - max_hsps 1 -num_threads 16 -evalue 1e-15”*. We only retained proteins with a significant BLAST hit for our final annotation, and used BUSCO, using the *Insecta* database, to assess annotation completeness.

### Mitochondrial genome assembly

We assembled and annotated the mitochondrial genome using Mitohifi (Allio et al. 2020; Uliano-Silva et al. 2021) on the scaffolded assembly, using the default parameters and the mitochondrial genome of *Anax parthenope* (Ma et al.) as a reference.

## Author Approvals

All authors have seen and approved this manuscript. It has not been submitted or published elsewhere.

## Acknowledgements

Hi-C data for the Black Petaltail were created by the DNA Zoo Consortium (dnazoo.org). DNA Zoo sequencing effort is supported by Illumina, Inc.; IBM; and the Pawsey Supercomputing Center. E.L.A. was supported by the Welch Foundation (Q-1866), a McNair Medical Institute Scholar Award, an NIH Encyclopedia of DNA Elements Mapping Center Award (UM1HG009375), a US-Israel Binational Science Foundation Award (2019276), the Behavioral Plasticity Research Institute (NSF DBI-2021795), NSF Physics Frontiers Center Award (NSF PHY-2019745), and an NIH CEGS (RM1HG011016-01A1). This work was also supported by a college undergraduate research award from the Department of Life Sciences at Brigham Young University.

## Data availability

The draft and unfiltered reference assembly are publicly available on DNA Zoo’s website (*https://www.dnazoo.org/assemblies/Tanypteryx_hageni (n.d.b)*). The filtered reference assembly is available on gennank. All genome annotation files can be found on figshare (https://figshare.com/projects/A_Chromosome-length_Assembly_of_the_Black_Petaltail_Tanypteryx_hageni_Dragonfly/151584).

## References

Aldous AR, Gannett MW, Keith M, O’Connor J. 2015. Geologic and Geomorphic Controls on the Occurrence of Fens in the Oregon Cascades and Implications for Vulnerability and Conservation. Wetlands. 35:757–767. doi: 10.1007/s13157-015-0667-x.

Allio R et al. 2020. MitoFinder: Efficient automated large-scale extraction of mitogenomic data in target enrichment phylogenomics. Molecular Ecology Resources. 20:892–905. doi: 10.1111/1755-0998.13160.

Baird I. 2012. The wetland habitats, biogeography and population dynamics of Petalura gigantea (Odonata: Petaluridae) in the Blue Mountains of New South Wales. PhD, The University of Western Sydney.

Bedford BL, Godwin KS. 2003. Fens of the United States: Distribution, characteristics, and scientific connection versus legal isolation. Wetlands. 23:608–629. doi: 10.1672/0277-5212(2003)023[0608:FOTUSD]2.0.CO;2.

Blanke A, Greve C, Mokso R, Beckmann F, Misof B. 2013. An updated phylogeny of Anisoptera including formal convergence analysis of morphological characters. Systematic Entomology. 38:474–490. doi: 10.1111/syen.12012.

Boulesteix M, Weiss M, Biémont C. 2006. Differences in Genome Size Between Closely Related Species: The Drosophila melanogaster Species Subgroup. Molecular Biology and Evolution. 23:162–167. doi: 10.1093/molbev/msj012.

Bybee SM, Ogden TH, Branham MA, Whiting MF. 2008. Molecules, morphology and fossils: a comprehensive approach to odonate phylogeny and the evolution of the odonate wing. Cladistics. 24:477–514.

Camacho C et al. 2009. BLAST+: architecture and applications. BMC Bioinformatics. 10:421. doi: 10.1186/1471-2105-10-421.

Campbell MS, Holt C, Moore B, Yandell M. 2014. Genome Annotation and Curation Using MAKER and MAKER-P. Current Protocols in Bioinformatics. 48:4.11.1-4.11.39. doi: 10.1002/0471250953.bi0411s48.

Cheng H, Concepcion GT, Feng X, Zhang H, Li H. 2021. Haplotype-resolved de novo assembly using phased assembly graphs with hifiasm. Nat Methods. 18:170–175. doi: 10.1038/s41592-020-01056-5.

Colles A, Liow LH, Prinzing A. 2009. Are specialists at risk under environmental change? Neoecological, paleoecological and phylogenetic approaches. Ecol Lett. 12:849–863. doi: 10.1111/j.1461-0248.2009.01336.x.

Danecek P et al. 2021. Twelve years of SAMtools and BCFtools. Gigascience. 10:giab008. doi: 10.1093/gigascience/giab008.

Dudchenko O et al. 2017. De novo assembly of the Aedes aegypti genome using Hi-C yields chromosome-length scaffolds. Science. 356:92–95. doi: 10.1126/science.aal3327.

Dudchenko O et al. 2018. The Juicebox Assembly Tools module facilitates de novo assembly of mammalian genomes with chromosome-length scaffolds for under $1000. doi: 10.1101/254797.

Durand NC et al. 2016. Juicer Provides a One-Click System for Analyzing Loop-Resolution Hi-C Experiments. cels. 3:95–98. doi: 10.1016/j.cels.2016.07.002.

Fiorillo AR. 2008. Dinosaurs of Alaska: Implications for the Cretaceous origin of Beringia. In: The Terrane Puzzle: New Perspectives on Paleontology and Stratigraphy from the North American Cordillera. Blodgett, RB & Stanley, GD, Jr, editors. Vol. 442 Geological Society of America p. 0. doi: 10.1130/2008.442(15).

Flynn JM et al. 2020. RepeatModeler2 for automated genomic discovery of transposable element families. Proceedings of the National Academy of Sciences. 117:9451–9457. doi: 10.1073/pnas.1921046117.

Fu L, Niu B, Zhu Z, Wu S, Li W. 2012. CD-HIT: accelerated for clustering the next-generation sequencing data. Bioinformatics. 28:3150–3152. doi: 10.1093/bioinformatics/bts565.

Galbraith DW, Lambert GM, Macas J, Dolezel J. 2001. Analysis of nuclear DNA content and ploidy in higher plants. Curr Protoc Cytom. Chapter 7:Unit 7.6. doi: 10.1002/0471142956.cy0706s02.

Hopkins MJ. 2011. How species longevity, intraspecific morphological variation, and geographic range size are related: a comparison using late Cambrian trilobites. Evolution. 65:3253–3273. doi: 10.1111/j.1558-5646.2011.01379.x.

Kohli M et al. 2021. Evolutionary history and divergence times of Odonata (dragonflies and damselflies) revealed through transcriptomics. iScience. 24:103324. doi: 10.1016/j.isci.2021.103324.

Laetsch DR, Blaxter ML. 2017. BlobTools: Interrogation of genome assemblies. doi: 10.12688/f1000research.12232.1.

Letsch H. 2007. Phylogeny of Anisoptera (Insecta: Odonata): promises and limitations of a new alignment approach. PhD Thesis, Rheinische Friedrich-Wilhelms-Universität in Bonn.

Li H. 2022. lh3/miniprot. https://github.com/lh3/miniprot (Accessed October 13, 2022).

Li H. 2018. Minimap2: pairwise alignment for nucleotide sequences. Bioinformatics. 34:3094– 3100. doi: 10.1093/bioinformatics/bty191.

Liu H et al. 2022. Chromosome-level genome of the globe skimmer dragonfly (Pantala flavescens). GigaScience. 11:giac009. doi: 10.1093/gigascience/giac009.

Ma F, Hu Y, Wu J. The complete mitochondrial genome of Anax parthenope (Odonata: Anisoptera) assembled from next-generation sequencing data. Mitochondrial DNA B Resour. 5:2817–2818. doi: 10.1080/23802359.2020.1789513.

Manni M, Berkeley MR, Seppey M, Simão FA, Zdobnov EM. 2021. BUSCO Update: Novel and Streamlined Workflows along with Broader and Deeper Phylogenetic Coverage for Scoring of Eukaryotic, Prokaryotic, and Viral Genomes. Molecular Biology and Evolution. 38:4647–4654. doi: 10.1093/molbev/msab199.

Misof B et al. 2014. Phylogenomics resolves the timing and pattern of insect evolution. Science. 346:763–767. doi: 10.1126/science.1257570.

Nardon C et al. 2005. Is genome size influenced by colonization of new environments in dipteran species? Mol Ecol. 14:869–878. doi: 10.1111/j.1365-294X.2005.02457.x.

Pertea G. 2022. gpertea/gffread. https://github.com/gpertea/gffread (Accessed October 13, 2022).

Pflug JM, Holmes VR, Burrus C, Johnston JS, Maddison DR. 2020. Measuring Genome Sizes Using Read-Depth, k-mers, and Flow Cytometry: Methodological Comparisons in Beetles (Coleoptera). G3 Genes|Genomes|Genetics. 10:3047–3060. doi: 10.1534/g3.120.401028.

Powell MG. 2007. Geographic Range and Genus Longevity of Late Paleozoic Brachiopods. Paleobiology. 33:530–546.

Price BW et al. 2022. The genome sequence of the blue-tailed damselfly, Ischnura elegans (Vander Linden, 1820). doi: 10.12688/wellcomeopenres.17691.1.

Ranallo-Benavidez TR, Jaron KS, Schatz MC. 2020. GenomeScope 2.0 and Smudgeplot for reference-free profiling of polyploid genomes. Nat Commun. 11:1432. doi: 10.1038/s41467-020-14998-3.

Rao SSP et al. 2014. A 3D Map of the Human Genome at Kilobase Resolution Reveals Principles of Chromatin Looping. Cell. 159:1665–1680. doi: 10.1016/j.cell.2014.11.021.

Robinson JT et al. 2018. Juicebox.js Provides a Cloud-Based Visualization System for Hi-C Data. cels. 6:256-258.e1. doi: 10.1016/j.cels.2018.01.001.

Shiryev SA, Papadopoulos JS, Schäffer AA, Agarwala R. 2007. Improved BLAST searches using longer words for protein seeding. Bioinformatics. 23:2949–2951. doi: 10.1093/bioinformatics/btm479.

Smit A, Hubley R, Green P. 2013. RepeatMasker Open-4.0. http://www.repeatmasker.org.

Stanke M, Diekhans M, Baertsch R, Haussler D. 2008. Using native and syntenically mapped cDNA alignments to improve de novo gene finding. Bioinformatics. 24:637–644. doi: 10.1093/bioinformatics/btn013.

Stanke M, Tzvetkova A, Morgenstern B. 2006. AUGUSTUS at EGASP: using EST, protein and genomic alignments for improved gene prediction in the human genome. Genome Biology. 7:S11. doi: 10.1186/gb-2006-7-s1-s11.

Suvorov A et al. 2021. Deep Ancestral Introgression Shapes Evolutionary History of Dragonflies and Damselflies. Systematic Biology. syab063. doi: 10.1093/sysbio/syab063.

Tolman DA. 2007. Soil Patterns in Three Darlingtonia Fens of Southwestern Oregon. naar. 27:374–384. doi: 10.3375/0885-8608(2007)27[374:SPITDF]2.0.CO;2.

Trizna M. 2020. assembly_stats 0.1.4. doi: 10.5281/zenodo.3968775.

Uliano-Silva M, Nunes JGF, Krasheninnikova K, McCarthy SA. 2021. marcelauliano/MitoHiFi: mitohifi_v2.0. doi: 10.5281/zenodo.5205678.

US Department of Agriculture. The Fen Phenomenon An Elusive Ecosystem. https://www.fs.usda.gov/Internet/FSE_DOCUMENTS/fseprd504763.pdf (Accessed April 14, 2022).

Ware JL et al. 2014. The petaltail dragonflies (Odonata: Petaluridae): Mesozoic habitat specialists that survive to the modern day. Journal of Biogeography. 41:1291–1300. doi: 10.1111/jbi.12273.

DNA Zoo. DNA Zoo. https://www.dnazoo.org (Accessed July 27, 2022a).

Tanypteryx_hageni. DNA Zoo. https://www.dnazoo.org/assemblies/Tanypteryx_hageni (Accessed July 27, 2022b).

