## Supplementary Materials for "A Chromosome-length Assembly of the Black Petaltail (*Tanypteryx hageni*) Dragonfly"

### *Atlantic Horseshoe Crab BLAST*

Using BLAST v 2.12.0, a *blastn* search was conducted using the *Limulus polyphemus* genome as a query against the standard NCBI BLAST database. This database currently includes the *L. polyphemus* genome, so the option *-negativetaxids* was included in the *blastn* search to exclude self-matching sequences. The resulting hits were then classified by phylum using NCBI's TaxonKit v 0.13.0 and visualized in RStudio using ggplot2 v 3.3.6.

### *Genome Size Estimation*

*D. melanogaster* heads and *T. hageni* head and thorax tissue were placed into ice cold Galbraith's buffer containing RNase A and crushed using a Dounce homogenizer. Lysate was then transferred to FACS tubes, through a filter. Samples were stained with propidium iodide (PI) diluted in Galbraith's buffer (Galbraith et al. 2001) (1mg/mL) for 3hrs. Two experiments were performed, one on the BD Accuri C6 cytometer and another on the Beckman Cytoflex cytometer, with two data points being retrieved from the Accuri experiment and one from the Cytoflex experiment. During the Accuri experiment, two data points were obtained from the ratios of red mean fluorescence intensities of the G1 and G2 peaks of one *Drosophila* sample and one *T. hageni* sample. During the cytoflex experiment, only the G1 peak of the *Drosophila* and *T. hageni* samples were compared. Thus, two biological replicates are presented with an additional technical replicate. The total quantity of DNA content within the *T. hageni* samples was calculated as the ratio of red MFI of the G1 and G2 peaks of the *T. hageni* samples to the G1 and G2 peaks of the *D. melanogaster* standards multiplied by the genome size estimates of *D. melanogaster*. As demonstrated by other insect genome estimates obtained by next-generation sequencing technology and flow cytometry, including *D. melanogaster*, flow cytometry analysis may result in larger genome size estimates compared to sequencing methods (Pflug et al. 2020).

**Supplementary References**

Supplementary references can be found in the main bibliography.

**Supplementary figure 1: GenomeScope Profile of the Black Petaltail**

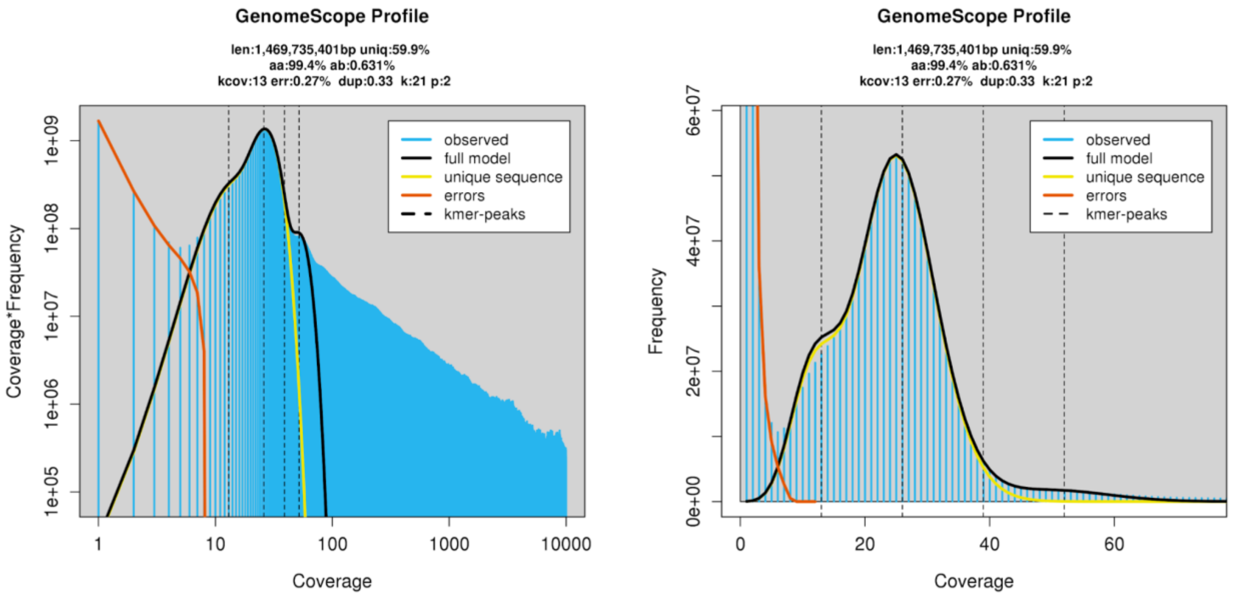


*Supplementary figure 1: The genome scope profile of the Black Petaltail, showing the relationships between coverage and frequency and coverage by coverage*frequency.*

**Supplementary Figure 2: Blobtools Output**

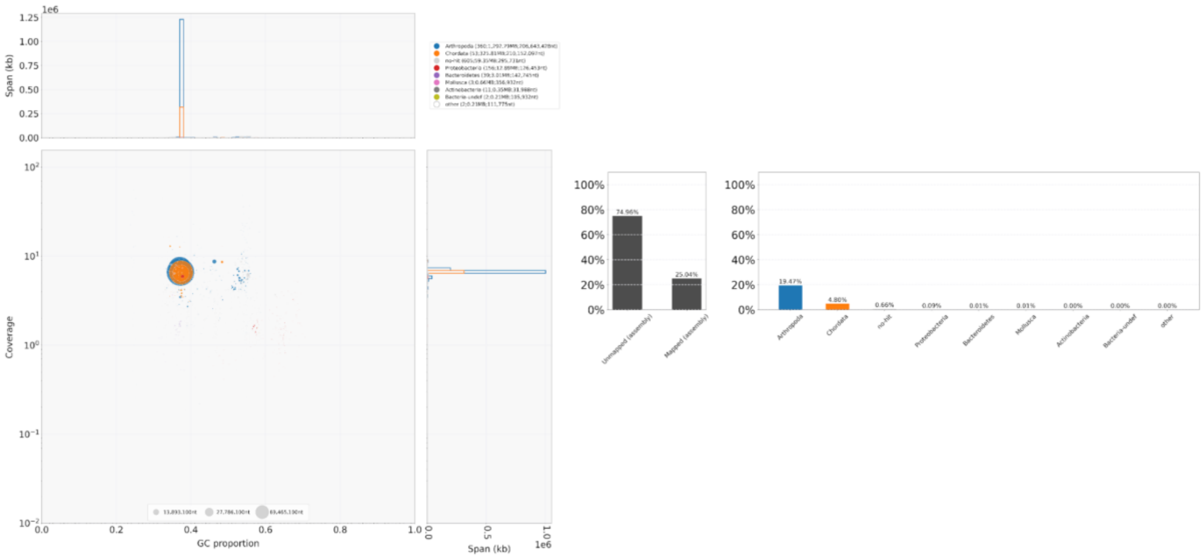


*Supplementary figure 2: Blobtools output of the Black Petaltail Genome. Note: contigs assigned to proteobacteria, bacteroides, mollusca and cnidaria were removed for the final reference assembly.*

**Supplementary figure 3: BLAST hits of Petaluridae Transcriptomes**

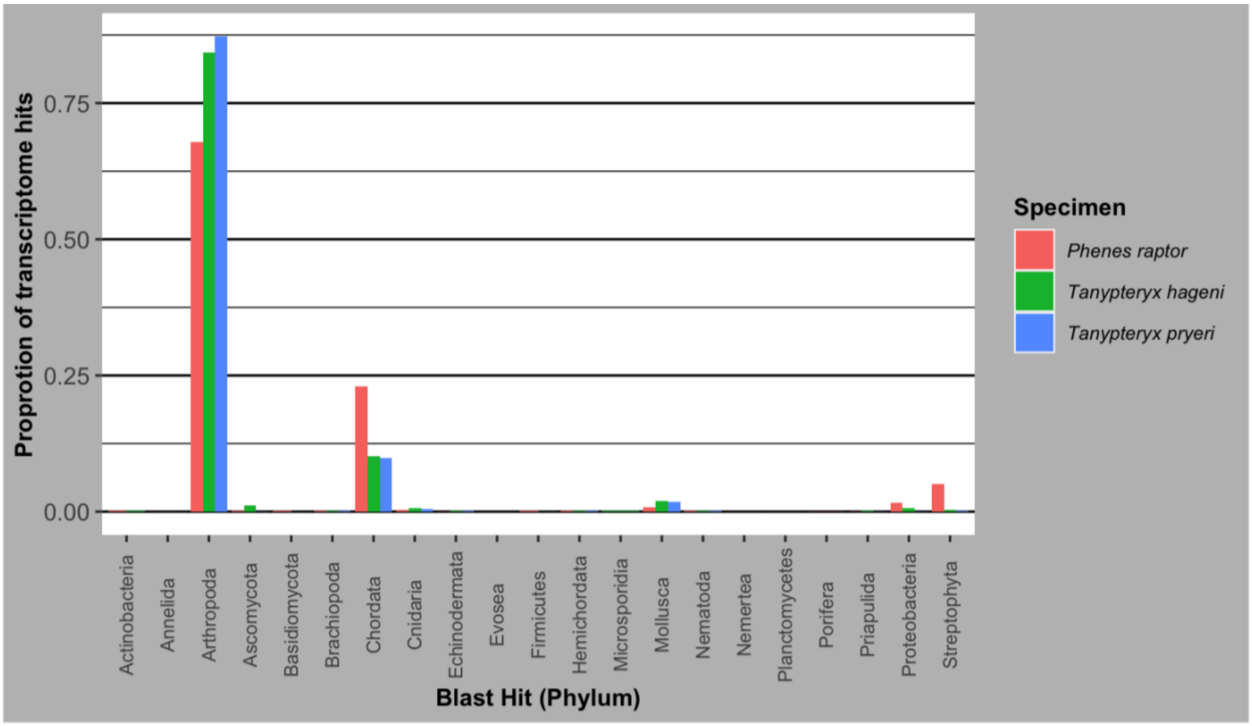


*Supplementary figure 3: Shows BLAST hits (by phylum) of the transcriptomes of Phenes raptor, Tanypteryx hageni, and Tanypteryx pryeri. More than 5% of all three transcriptomes blasted to chordata.*

**Supplementary figure 4: BLAST hits of Atlantic Horseshoe Crab genome assembly**

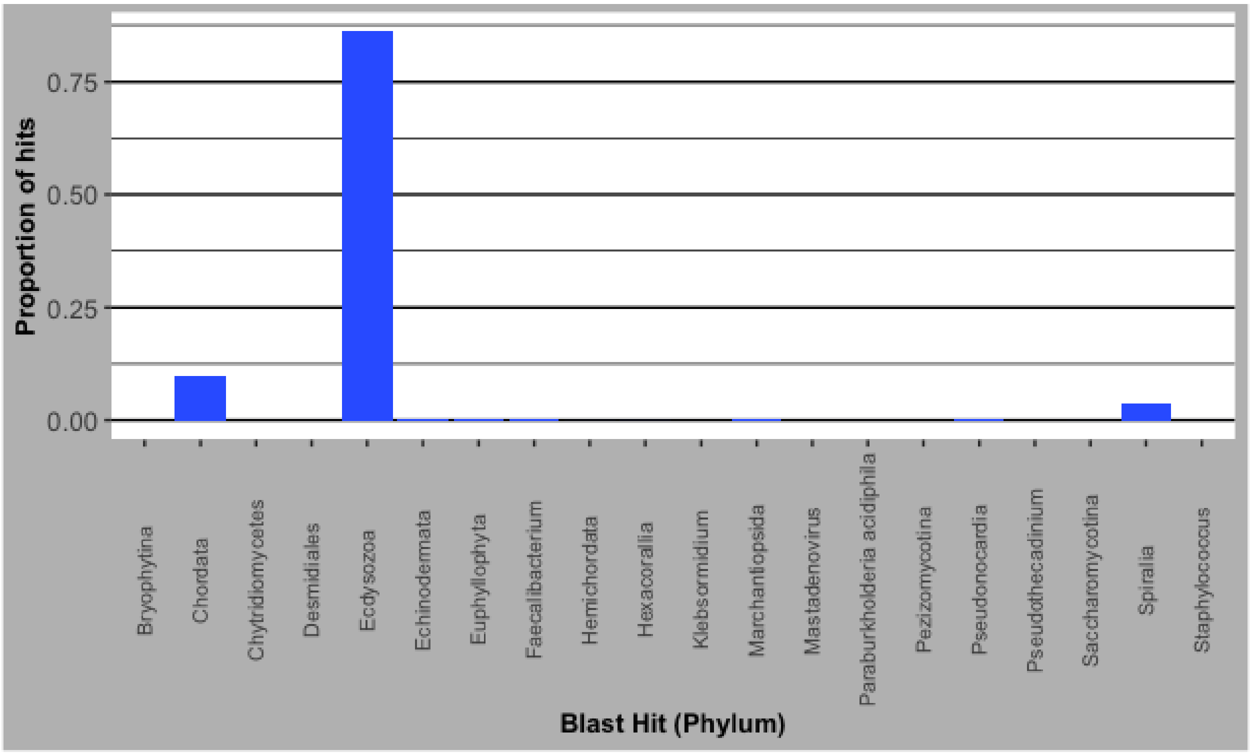


*Supplementary figure 4: BLAST hits (by phylum) of the Limulus polyphemus genome. Chordata accounted for 9.7% of the total hits.*

**Supplementary figure 5: Interactive contact map of the chromosome length assembly**

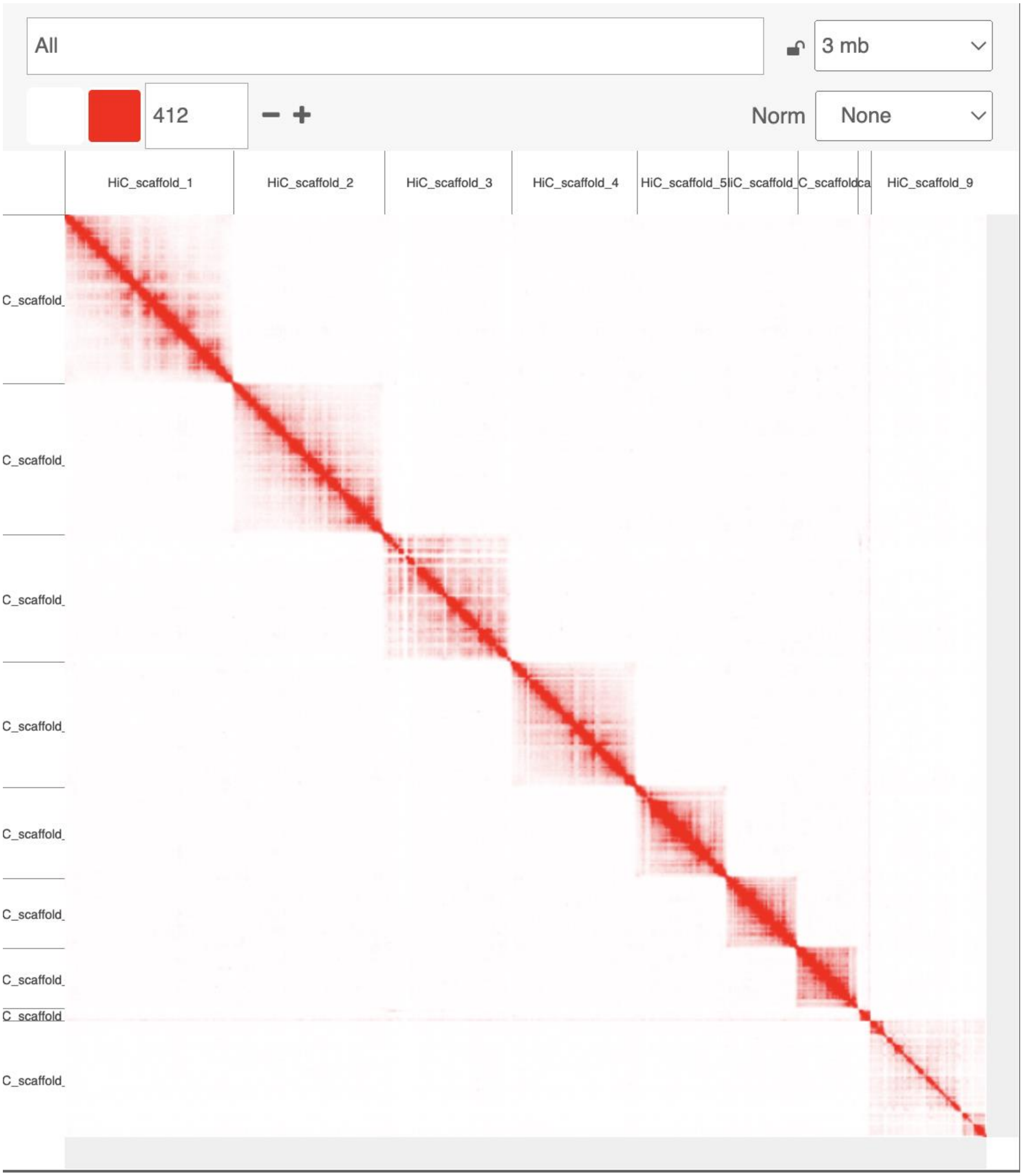


*Supplementary figure 5: Interactive contact map of the chromosome-length assembly with Hi-C scaffolding.*

**Supplemental figure 6: Flow cytometry estimates**

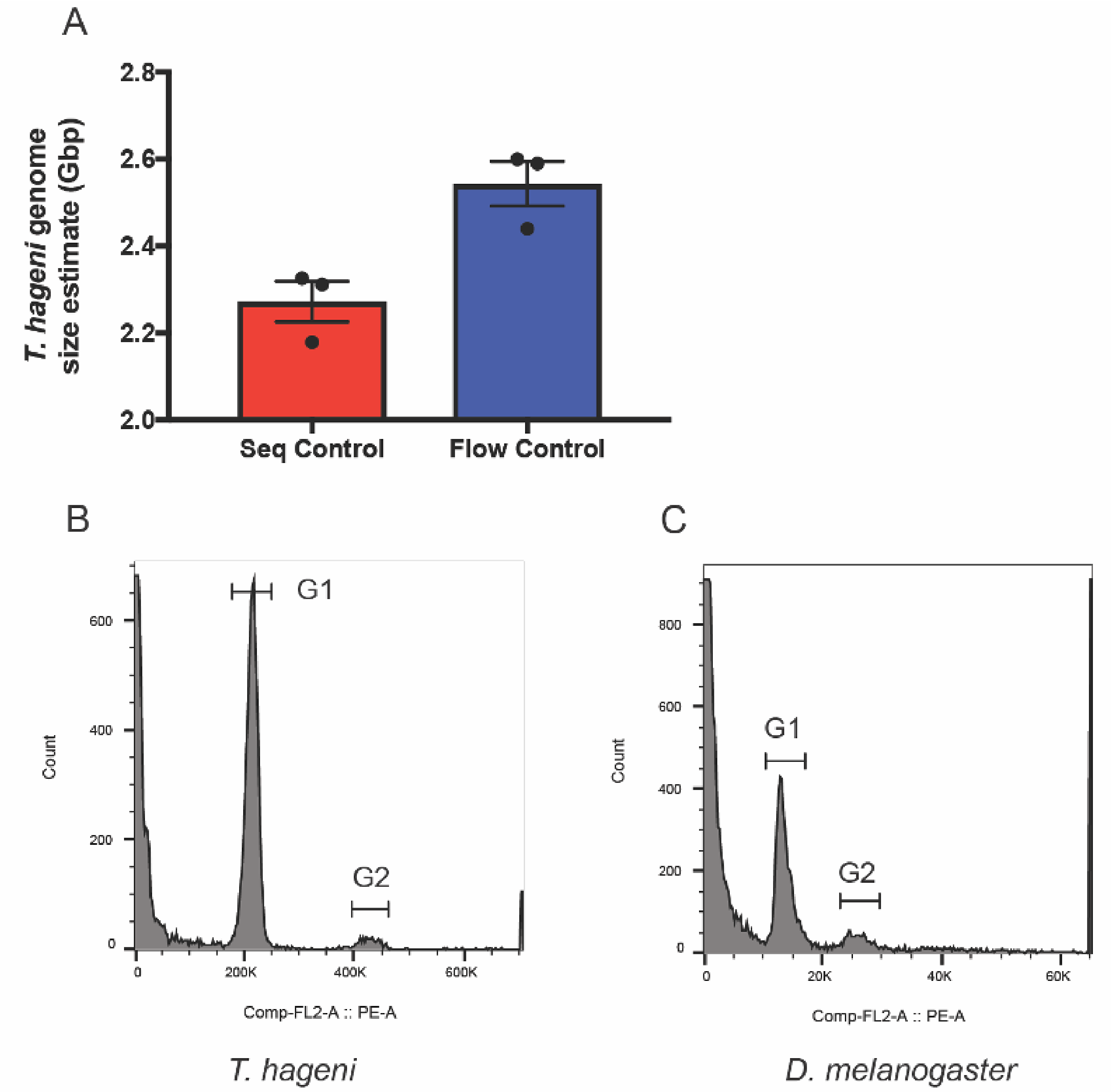


**Supplemental Figure 6. *Tanypteryx hageni* genome size estimate using flow cytometry.** (A) Two *Tanypteryx hageni* genome size estimates calculated using flow cytometry. The red bar is the *T. hageni* genome estimate (2.26 Gbp) when calculated using the *D. melanogaster* sequenced genome (mean size of 138.9 Mbp) as the control. The blue bar is the *T. hageni* genome estimate (2.54 Gbp) when calculated using the low flow cytometry estimate of the *D. melanogaster* genome (156 Mbp) as the control. (B) Representative flow cytometry histogram of *T. hageni* head and thorax tissue stained with propidium iodide and gates added to G1 and G2 peaks. (C) Representative flow cytometry histogram of *D. melanogaster* head tissue stained with propidium iodide and gates added to G1 and G2 peaks.

| Supplementary Table I: Assembly statistics of iterations of the Black Petaltail genome | | | |
|---|---|---|---|
| Assembly | Number of Contigs/Scaffolds | N50 (mb) | Assembly Size (gb) |
| Draft | 2,133 | 4.2 | 1.69 |
| Unfiltered Chromosome Length Assembly | 1,231 | 206.6 | 1.70 |
| Filtered Chromosome Length Assembly | 1,033 | 206.6 | 1.68 |

*Supplementary table 1: Displays the assembly statistics of the draft, unfiltered chromosome length, and filtered chromosome length assemblies.*

| **Supplementary Table II: Chromosomes and Organelles of the Genome Assembly** | | |
|---|---|---|
| Contig | GC content | Size(bp) |
| Chromosome 1 | 39.15% | 353,118,706 |
| Chromosome 2 | 38.23% | 267,812,038 |
| Chromosome 3 | 38.16% | 236,878,591 |
| Chromosome 4 | 37.29% | 210,050,401 |
| Chromosome 5 | 37.25% | 152,463,193 |
| Chromosome 6 | 37.60% | 117,929,689 |
| Chromosome 7 | 37.41% | 105,601,530 |
| Chromosome 8 | 37.90% | 35,320,507 |
| Chromosome 9 | 37.18% | 198,988,126 |
| Mitochondrion | 24.62% | 16,053 |
| *Supplementary Table 2: Compares GC content and size of each chromosome and the mitochondrion in the genome assembly of the Black Petaltail.* | | |
